# Structure can emerge from disorder under neutral evolution

**DOI:** 10.64898/2026.05.28.728420

**Authors:** Bharat Ravi Iyengar, Erich Bornberg-Bauer

**Affiliations:** Institute for Evolution and Biodiversity, University of Münster, Hüfferstraße 1, 48149 Münster, Germany

## Abstract

Mutations are often thought to destabilize protein structures. However, many proteins are naturally unstructured and the effect of mutations on such proteins especially in the absence of selection, remains unclear. Here, we develop a computational model to study the effects of mutations on structural disorder in both random and natural sequences, including those derived from evolutionary conserved proteins, and evolutionarily novel *de novo* proteins.

We find that while structured proteins tend to lose structure, unstructured proteins exhibit the opposite trend, becoming more structured in the absence of directional selection. This bidirectional dynamics is robust to mutation biases, genetic code structure, and sequence origin, suggesting that it arises from the topology of the sequence–structure landscape rather than intrinsic directional mechanisms. Our results are consistent with a diffuse landscape in which structured and disordered sequences are interspersed throughout sequence space.

These findings suggest that neutral evolution can result in structure formation and may facilitate the early structural evolution of *de novo* proteins prior to strong selection.

## Introduction

Proteins often require a well-defined structure to perform their biological functions, as reflected by the prevalence of structured conformations among physiologically important proteins. However, numerous intrinsically disordered proteins (IDPs) also fulfill essential biological roles (van der Lee *et al*., 2014; Wright and Dyson, 2014; Holehouse and Kragelund, 2023). For example, intrinsically disordered regions can act as flexible linkers that tether catalytic kinase domains to their target proteins (Dyla and Kjaergaard, 2020). Disordered proteins can also form high-affinity complexes in the absence of well-defined structured interaction domains (Borgia *et al*., 2018). Thus, both structured and disordered proteins can be functional and maintained by selection (Zarin *et al*., 2019).

For structured proteins, purifying selection is expected to be strong because most mutations are destabilizing (Eyre-Walker and Keightley, 2007). This is consistent with the idea that a stable fold requires a precise arrangement of physicochemical interactions and stereochemical constraints, and that mutations tend to perturb this optimized configuration. At the same time, some proteins such as ribosomal proteins, may exhibit substantial mutational robustness owing to highly stable structural interactions (Lind *et al*., 2010). In contrast, a comparable understanding of how mutations affect the structural properties of disordered proteins remains limited. More broadly, extant structured proteins represent only a minute fraction of all possible sequences, and the sequence–structure relationship, including mutational effects, remains largely unexplored across most of sequence space.

A sequence space is defined as the set of all possible protein sequences. Mapping sequences to their structural properties gives rise to a sequence–structure landscape. If stable structures require specific sequence compositions, then structured sequences should be relatively rare and confined to localized regions of this space. Moreover, because evolution usually proceeds through descent, many known proteins are expected to occupy proximal regions of sequence space. Together, these considerations suggest a landscape that is predominantly flat (corresponding to disordered sequences), with structured peaks confined to localized and potentially clustered regions. In such a scenario, the emergence of structured sequences through neutral drift alone is expected to be unlikely.

Several studies have explored the navigability of the sequence–structure landscape. Work on the distribution of fitness effects (DFE) of mutations has provided insights into the local structure of the landscape around specific proteins, showing that most mutations are deleterious and tend to destabilize structure (Eyre-Walker and Keightley, 2007; Wylie and Shakhnovich, 2011; Boucher *et al*., 2016). More detailed studies have examined navigability in local fitness landscapes that are rugged, where multiple optimal structures are separated by suboptimal configurations, often referred to as fitness valleys – regions of sequence space corresponding to low-fitness or unstable states (Papkou *et al*., 2023; Schulz *et al*., 2025). Despite this ruggedness, these studies demonstrate that optimal states can be reached under selection. Complementary approaches using simplified protein models, such as lattice models, have investigated landscape properties more generally through simulations (Govindarajan and Goldstein, 1997; Bastolla *et al*., 1999; Bornberg-Bauer and Chan, 1999; Greenbury *et al*., 2022). These studies suggest that stable configurations can be accessed from diverse initial conditions, partly because deep fitness valleys are relatively rare, facilitating movement between optima under selection.

Taken together, existing work indicates that selection can enable proteins to attain structured states, even when starting from disordered configurations or navigating rugged landscapes. However, these studies primarily focus on local regions of sequence space or rely on sim-plified models. As a result, the global properties of the sequence–structure landscape — including its topology and navigability across the vast space of previously uncharacterized sequences — remain poorly understood. In particular, it is unclear how neutral evolution shapes structural properties, and whether it can promote structure formation in disordered sequences while destabilizing structured ones.

To address these questions, we developed a computational model to study the structural evolution of both random protein sequences and biologically derived sequences, including *de novo* and conserved proteins, under neutral evolution. Specifically, we subjected each sequence to 30 generations of simulated evolution to explore the local sequence space and quantified structural disorder at each generation to map the sequence–structure landscape. Our simulations reveal that disordered sequences tend to become more structured under neutrality, while structured sequences tend to lose structure. Notably, this bidirectional change does not arise from directional biases inherent to the genetic code or mutation rates. Together, these findings demonstrate that neutral evolution can be constructive, enabling the emergence of structure from disorder. Furthermore, the presence of directional change in the absence of explicit directional mechanisms suggests that the underlying sequence– structure landscape is diffuse, with structured sequences interspersed among disordered ones.

## Results

### Model design

To understand how mutations affect the structure of non-adapted proteins, we developed and simulated a computational evolutionary model (Figure 1). To this end, we first generated random protein sequences, which we reasoned to be suitable proxies for non-adapted proteins. Specifically, we generated sequences parameterized by sequence length and amino acid frequencies (composition).

**Figure 1:**
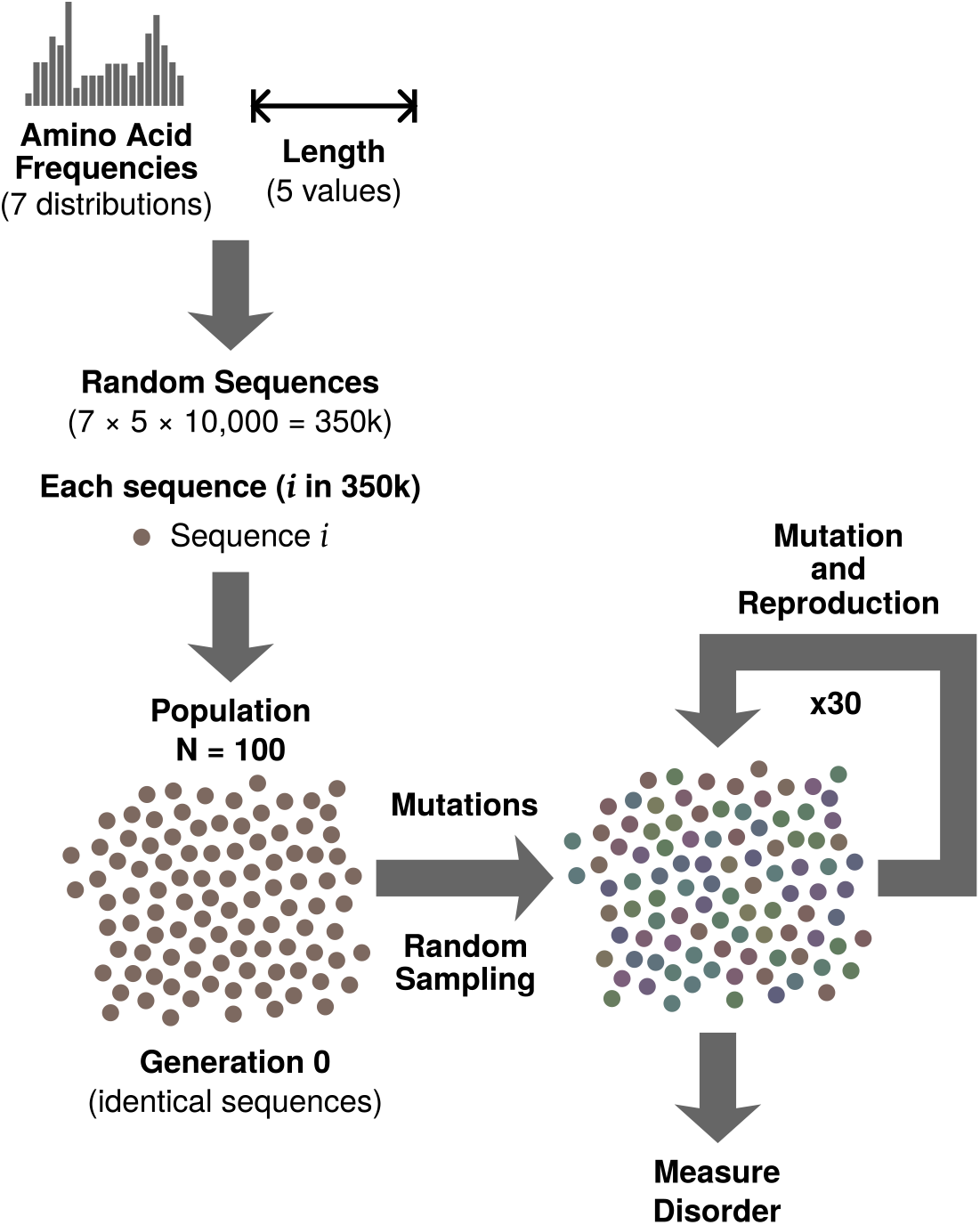
Overview of the simulation framework. Random protein sequences are generated across combinations of amino acid frequency distributions and sequence lengths. Each sequence initializes a population of identical sequences (*N* = 100), which evolves through repeated cycles of mutation and random sampling for 30 generations. Structural disorder is quantified for the evolved protein variants across generations.

We considered five sequence lengths (10, 30, 60, 90, and 120) and seven amino acid frequency distributions. The first six were defined by GC (guanine–cytosine) content levels of 30%, 35%, 40%, 45%, 50%, and 55%, while the seventh corresponded to a uniform distribution. For each combination of length and composition, we generated 10,000 sequences, resulting in a total of 350,000 sequences.

Next, we developed a stochastic model to simulate protein evolution, based on the Wright – Fisher model (Hartl and Clark, 1997). Each simulation begins with a population of 100 identical sequences (generation 0). At each subsequent generation, sequences may undergo mutation and reproduction. We modeled the number of mutations per sequence as a Poisson-distributed random variable with mean 1, allowing for zero or multiple mutations per sequence. Mutations occur uniformly along the sequence, and the type of mutation is deter-mined by an amino acid substitution matrix. In our primary simulations, we generated the substitution matrix using nucleic acid mutation biases reported in *Drosophila melanogaster* (Schrider *et al*., 2013; Iyengar and Bornberg-Bauer, 2023), although the results are qualitatively insensitive to the choice of mutation bias.

Reproduction occurs via sampling with replacement, leading to non-overlapping generations. To model neutral evolution, we did not impose any selection and instead tracked the phenotype (structural disorder) of the evolving sequences.

To quantify structural disorder, we used the computational disorder predictor ADOPT (Redl *et al*., 2023). We chose this tool for two reasons. First, it captures the global context of protein sequences using the ESM-1b protein language model. Second, it is trained on NMR-based structural data, making it suitable for previously uncharacterized sequences, unlike meth-ods trained on databases of proteins with annotated intrinsically disordered regions. We assigned a single representative score (*Z*) to each protein sequence, where higher values indicate lower levels of structural disorder, and *vice versa*. We also corroborated our results using a conceptually different disorder predictor IUPred3 (Erdős *et al*., 2021).

We subjected each of the 350,000 random sequences to evolutionary simulations for 30 generations. This relatively short timescale allows us to map the local mutational effect landscape, which is the primary goal of our study.

### Neutral mutations drive disordered proteins toward structure and structured proteins toward disorder

We hypothesized that, while random mutations are likely to disrupt the structure of highly structured proteins, they may have limited phenotypic effects on intrinsically disordered proteins. To test this hypothesis, we analyzed the relationship between *Z* scores and time (in generations) across evolutionary trajectories by computing the Spearman rank correlation between the two quantities. While a small fraction of trajectories showed statistically insignificant correlations (consistent with the null hypothesis), the majority (*>*80%) showed significant correlations between *Z* score and time (significance threshold *α* = 10^*−*3^). Interestingly, many trajectories (*>*75% in some length–composition combinations) showed a positive correlation, suggesting that protein sequences can become more structured through neutral drift alone (Figure 2A).

**Figure 2:**
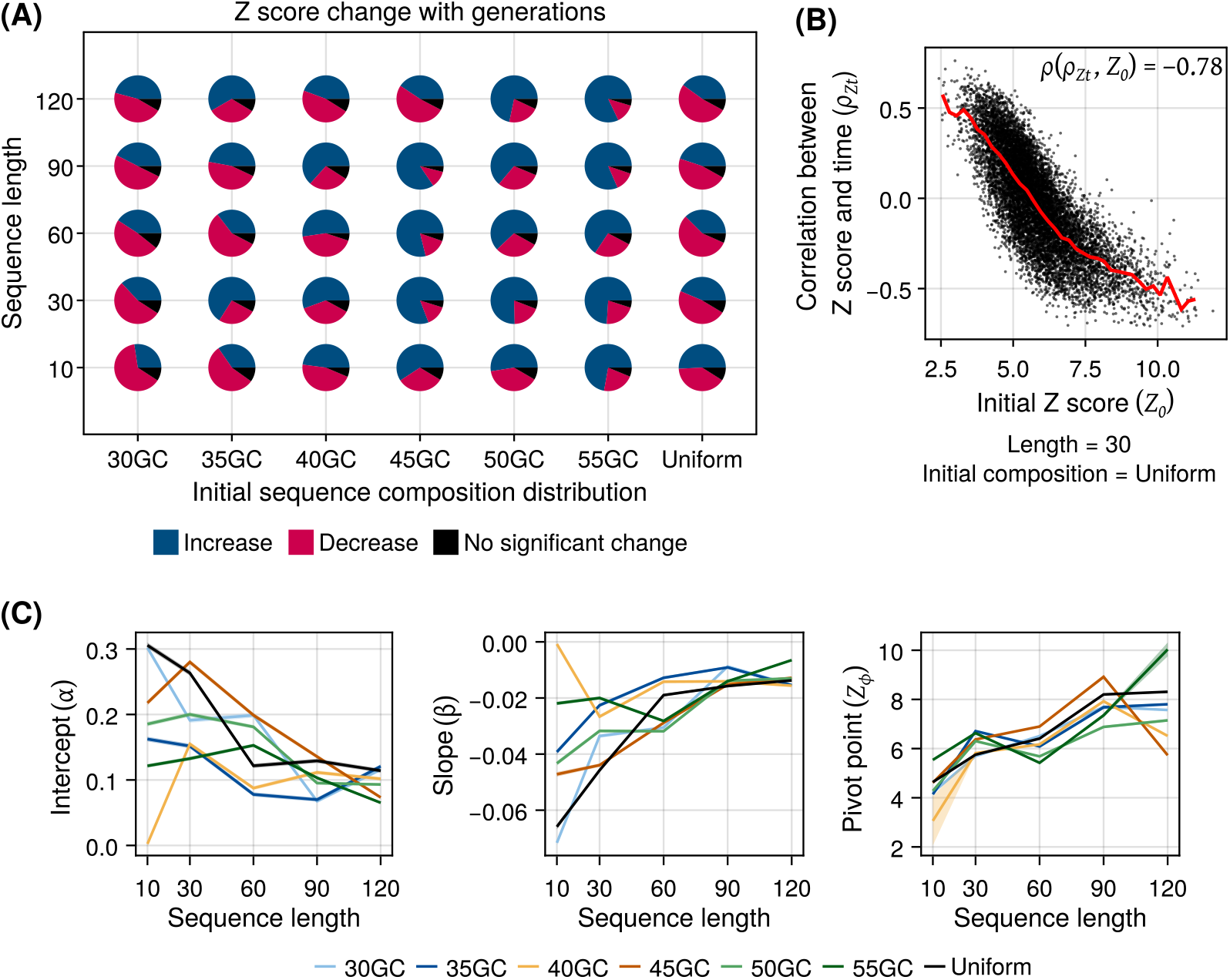
Neutral mutations drive opposing changes in protein disorder depending on initial state. **(A)** Pie charts showing the fraction of simulated evolutionary trajectories (out of 10,000) that exhibit an increase (blue), decrease (red), or no significant change in the *Z* score over time, across different combinations of sequence length (vertical axis) and initial composition (horizontal axis). Statistical significance is assessed using a Spearman correlation test. **(B)** Scatter plot showing the relationship between temporal changes in *Z* (quantified by *ρ*_*Zt*_; vertical axis) and the initial *Z* score (*Z*_0_; horizontal axis), for trajectories with sequence length 30 and uniform initial composition. The fitted trend line is shown in red. The inset reports the Spearman correlation between *ρ*_*Zt*_ and *Z*_0_. For other length–composition combinations, see Figure S1. **(C)** Fitted coefficients (*α* and *β*) and corresponding pivot points (*Z*_*ϕ*_; vertical axes) from the linear model, plotted as a function of sequence lengths (horizontal axes) and initial compositions (line colors). Shaded ribbons indicate 95% confidence intervals for the fitted values

Next, we investigated the factors determining the direction of disorder change. Based on our assumption that mutations increase disorder in structured proteins, we analysed the relationship between the initial *Z* score (*Z*_0_) and the correlation coefficient between *Z* and time (*ρ*_*Zt*_). We found that high values of *Z*_0_ are indeed associated with negative correlations (*ρ*_*Zt*_ *<* 0). More interestingly, as *Z*_0_ decreases, *ρ*_*Zt*_ increases, transitioning from negative to positive values and continuing to increase in magnitude (Figure 2B, Figure S1). Furthermore, this negative correlation between *Z*_0_ and *ρ*_*Zt*_ is statistically significant (t-test *P <* 10^*−*99^). To ensure that this pattern is not a technical artifact of the ADOPT algorithm, we performed the same analysis using disorder scores predicted by IUPred3 and obtained consistent results (Figure S2). This confirms that the inverse relationship between initial disorder and disorder change is robust to the choice of disorder predictor. Together, these results show that mutations can both disrupt structure in highly structured proteins and promote structure formation in disordered proteins.

We used a linear model to analyse this observation more formally. To this end, we developed a model that aligns mechanistically with the evolutionary simulations and respects the Markov property of the stochastic process. Specifically, we defined the mean change in *Z* score (Δ*Z*_*t*_) as a linear function of the mean population *Z* score (*Z*_*t*_) at a given time point (Equation 1). This formulation implicitly captures the effect of the initial *Z* score on Δ*Z*, while generalizing the relationship to all time points.

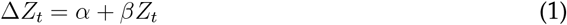

We found that the coefficient *β* is negative across all length–composition combinations, indicating that Δ*Z*_*t*_ decreases with increasing *Z*_*t*_. The coefficient *α*, which describes the baseline value of Δ*Z*_*t*_ when the *Z* score is (hypothetically) zero, is consistently positive, reinforcing the notion that highly disordered sequences tend to become less disordered under neutral drift (Figure 2C). Both *α* and *β* show greater variability across initial compositions for shorter sequences, suggesting that while disorder changes in short sequences are qualitatively pre-dictable, they remain sensitive to sequence composition.

A more informative descriptor of the effect of *Z*_0_ on Δ*Z*_*t*_ is the pivot point at which Δ*Z*_*t*_ transitions from positive to negative. This pivot point (*Z*_*ϕ*_), defined by Δ*Z*_*t*_ = 0, is given by *−α/β*. Notably, *Z*_*ϕ*_ is more robust to variation in initial composition than the individual coefficients, indicating that short-term evolutionary trajectories are governed by a characteristic balance point despite variability in the underlying parameters (Figure 2C). However, *Z*_*ϕ*_ varies more strongly with sequence length than with initial composition, showing an apparent increase with increasing length. This relationship is not monotonic and cannot be fully captured by a linear interaction model. However, it illustrates an overall trend where longer sequences tend to exhibit a positive Δ*Z*_*t*_ under neutral evolution, consistent with a higher *Z*_*ϕ*_, while shorter sequences with a comparable *Z*_*t*_ tend to exhibit a negative Δ*Z*_*t*_, consistent with a lower *Z*_*ϕ*_.

### Directional disorder changes are not driven by mutation bias or the structure of the genetic code

Next, we investigated the mechanisms that could explain directional changes (Δ*Z*) arising from non-directional processes such as neutral drift. Although mutations occur randomly, their probabilities are shaped by mutation rate biases and the structure of the genetic code. First, amino acids with more degenerate codons are more likely to accumulate over time. Second, the genetic code constrains which amino acid substitutions are more likely; for example, a glycine-to-alanine substitution requires a single nucleotide change, whereas a glycine-to-proline substitution requires at least two. Third, different nucleotide mutations occur at different rates, with transitions (e.g., A→G, G→A) generally more frequent than transversions (e.g., A→C, G→T). Together, these factors can introduce directionality into amino acid substitutions (Iyengar and Bornberg-Bauer, 2023).

To test whether such biases drive the observed changes in disorder, we performed simulations in which these factors were randomized. Specifically, we repeatedly generated randomized amino acid substitution matrices using the randomization procedure and performed independent simulations using a unique matrix for each trajectory.

We first examined the effect of mutation bias while keeping the genetic code structure intact but varying nucleotide substitution rates. Fitting the linear model described in Equation 1, we estimated the coefficients (*α* and *β*) and the pivot point (*Z*_*ϕ*_). Across all length–composition combinations, *α* remained positive and *β* remained negative, consistent with our earlier observation that disordered sequences tend to become more structured and structured sequences tend to lose order. We also observed a non-monotonic increase in *Z*_*ϕ*_ with sequence length. These results indicate that biased nucleotide mutation rates do not explain the directionality of disorder changes (Figure 3).

**Figure 3:**
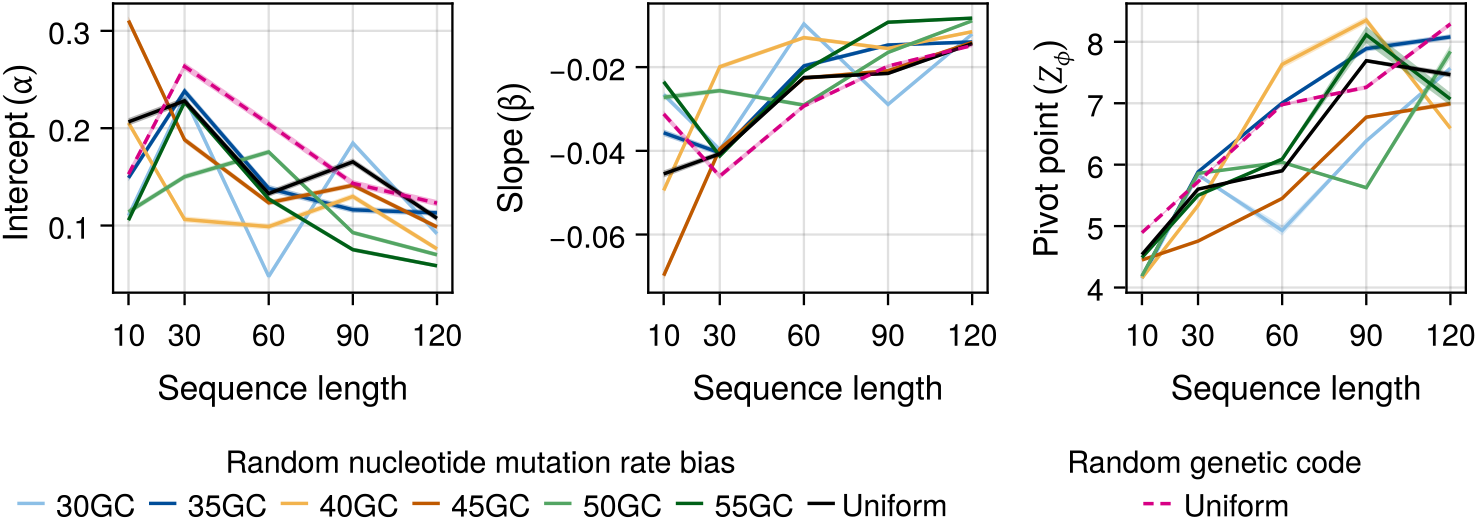
Fitted coefficients (*α* and *β*) and corresponding pivot points (*Z*_*ϕ*_; vertical axes) from the linear model, plotted as a function of sequence lengths (horizontal axes) and initial compositions (line colors). Coefficient values and corresponding pivot points estimated from trajectories generated under randomized mutation bias and randomized genetic code are indicated by solid and dashed lines, respectively. Shaded ribbons indicate 95% confidence intervals for the fitted values.

We next examined the effect of genetic code structure, which can impose stronger constraints on amino acid substitutions. To this end, we simulated the evolution of random sequences with a uniform initial amino acid distribution, as GC content cannot be mapped to amino acid frequencies under a randomized genetic code. As before, we observed a positive *α* and a negative *β* across all sequence lengths, along with a length-dependent increase in *Z*_*ϕ*_ (Figure 3).

Overall, both the simulations randomizing mutation bias and the genetic code structure yield qualitatively similar results to the original model. These findings indicate that the observed directional changes in disorder are not driven by specific mutational biases or features of the genetic code, but instead reflect intrinsic properties of the sequence–structure landscape.

### Directional disorder changes in natural sequences is consistent with that of random sequences

Next, we investigated if naturally occurring protein sequences also undergo directional changes in their structure under neutral evolution, like random sequences. To this end, we analysed two classes of sequences compiled in a previous study (Heames *et al*., 2020) – *de novo* and conserved proteins with a comparable length distribution (22 – 250 amino acids). While *de novo* sequences are not completely random, they are evolutionarily novel and lack detectable homology to conserved proteins. In contrast, the conserved proteins in this dataset include both structured and disordered sequences, providing a diverse range of initial structural states to test our model.

We subjected 2269 conserved and 2381 *de novo* sequences to simulated neutral evolution and analysed the changes in *Z* score over time. Evolutionary trajectories of both sequences classes exhibited qualitatively similar behavior to that of random sequences. Both sequence classes experienced bidirectional changes in their *Z* score (Figure 4A) where the correlation between *Z* score and time (*ρ*_*Zt*_) depended negatively with the initial *Z* score (*Z*_0_), with *ρ*_*Zt*_ *<* 0 for higher values of *Z*_0_ and *ρ*_*Zt*_ *>* 0 for lower values of *Z*_0_ (Figure 4B). Fitting the linear model to these trajectories (Equation 1) yielded a positive *α* and a negative *β* for both sequence classes, consistent with a decrease in Δ*Z*_*t*_ with increasing *Z*_*t*_ (Figure 4C). However, the pivot point of conserved sequences (~7.8) was higher than that of the *de novo* sequences (~5.7) indicating that neutral evolution tends to reduce disorder in conserved sequences at a broader range of *Z* scores.

**Figure 4:**
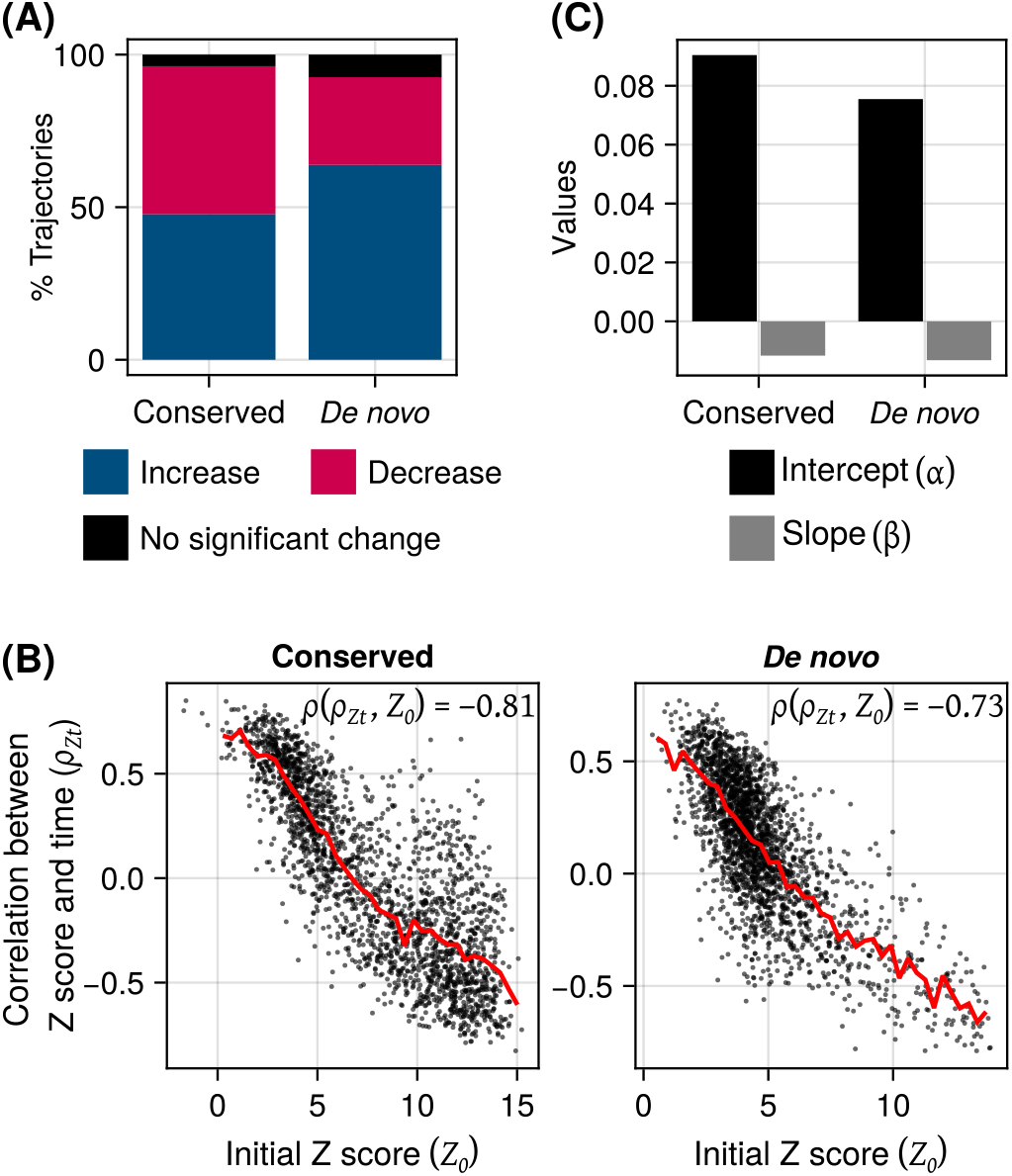
Bidirectional disorder changes in natural (conserved and *de novo*) sequences. **(A)** Stacked bars denoting the percentage of trajectories that exhibited positive (blue), negative (red) or non-significant changes (black) in *Z* score over time (Spearman correlation test), for conserved and *de novo* sequences (horizontal axis). **(B)** Scatter plots showing the relationship between temporal changes in *Z* (quantified by *ρ*_*Zt*_; vertical axes) and the initial *Z* score (*Z*_0_; horizontal axes), for conserved and *de novo* sequences. The insets report the Spearman correlation between *ρ*_*Zt*_ and *Z*_0_. **(C)** Grouped bars showing the values (vertical axis) of fitted the coefficients – *α* (black) and *β* (gray) from the linear model (Equation 1), for conserved and *de novo* sequences (horizontal axis). The pivot estimates (*Z*_*ϕ*_) for the conserved and *de novo* sequences are ~7.8 and ~5.7, respectively.

**Figure 5:**
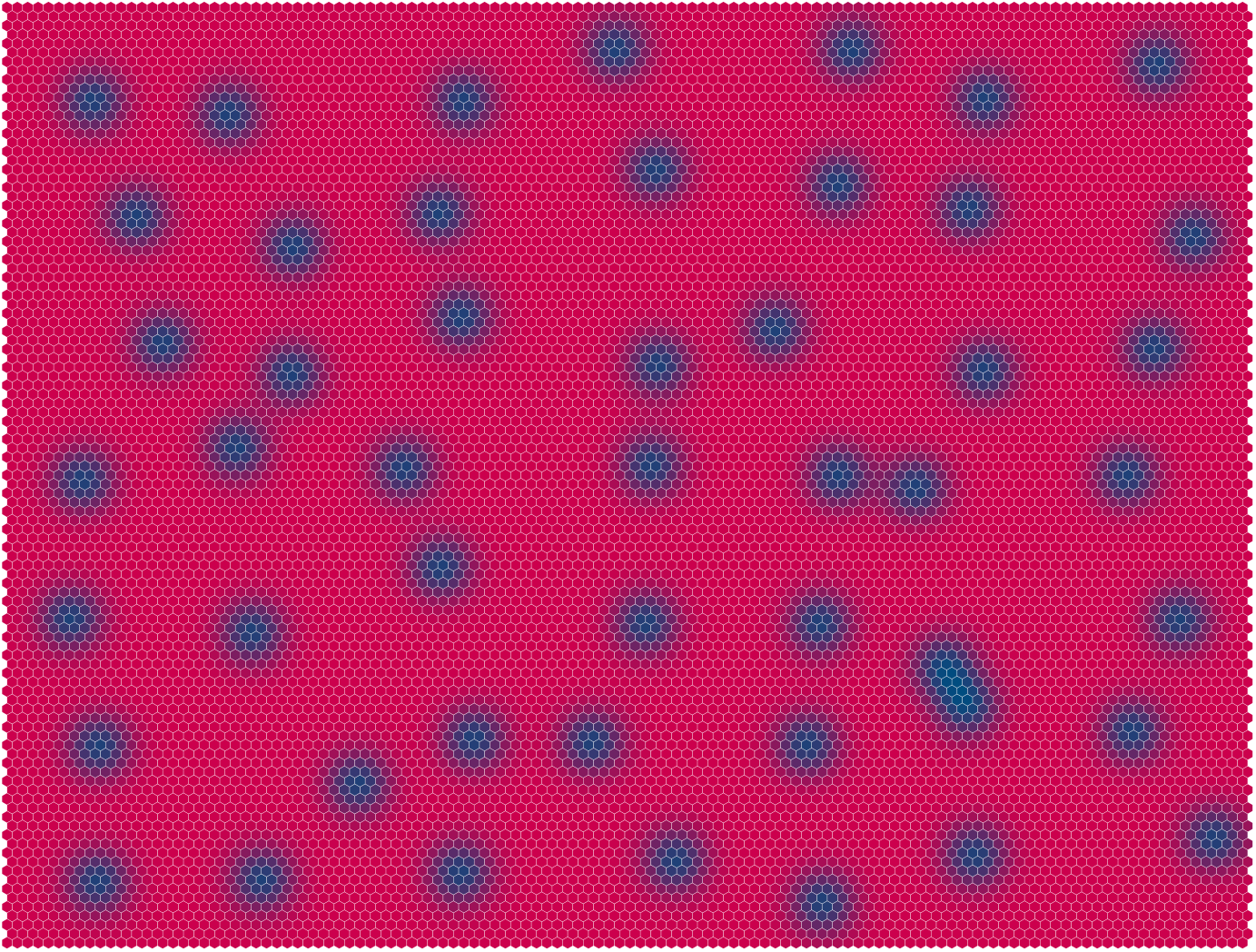
A schematic representation of the sequence–structure landscape suggested by the model results, illustrated as a hexagonal honeycomb grid in which similar sequences occupy neighboring cells. Cell colors denote the structural disorder of the corresponding sequences, with red and blue representing high and low disorder respectively. The figure illustrates how local neutral mutational trajectories (random walks) originating from highly disordered sequences (red) can lead to increased structural order (blue).

Overall, these results show that the directional changes in structural disorder observed in random sequences are also present in biologically derived protein sequences, indicating that the underlying sequence–structure relationships captured by our model extend beyond purely random sequence space.

## Discussion

A longstanding question in protein science is how genotype–phenotype relationships are distributed across sequence space – specifically, whether folded and disordered proteins occupy distinct regions or are interspersed throughout. While it is infeasible to survey large portions of sequence space directly, it is possible to explore the local landscape around a diverse set of focal sequences.

Here, we performed such an exploration using 350,000 focal protein sequences spanning a range of lengths and amino acid compositions, with the goal of characterizing the sequence–structure relationship. We selected these sequences by random sampling to obtain an unbiased representation of sequence space. This choice is motivated by three considerations. First, random sequences provide access to a broad diversity of phenotypes. Second, they serve as proxies for non-adapted proteins, allowing us to probe regions of sequence space that are largely unexplored. Finally, naturally occurring proteins may occupy restricted regions of sequence space due to shared evolutionary ancestry, which could bias inferences about the global distribution of sequence–structure relationships.

We addressed this question by simulating the evolution of the focal sequences and measuring structural disorder at each step. Importantly, we imposed no selection on structure, allowing evolutionary trajectories to explore a broader region of sequence space, in contrast to directed evolution, which constrains exploration to sequences optimized for foldability. We focused on relatively short protein sequences (10 – 120 amino acids) for both biological and methodological reasons. Biologically, short sequences are more likely to arise *de novo* (Bornberg-Bauer *et al*., 2021; Iyengar and Bornberg-Bauer, 2023), and thus serve as plausible models of non-adapted proteins. Methodologically, shorter sequences allow efficient exploration of the local mutational landscape, as a modest number of mutations can substantially diversify the sequence while retaining proximity to the initial state. Accordingly, we limited the simulations to 30 generations, corresponding to an average of 30 mutations per sequence. This timescale is sufficient to diversify short proteins while still restricting exploration to the local neighborhood of the initial sequences. Together, this design enables controlled probing of genotype – phenotype relationships without conflating effects from more distant regions of sequence space.

Our simulations show that structured proteins tend to become more disordered under mutations, consistent with the well-established destabilizing effects of mutations in conserved proteins (Eyre-Walker and Keightley, 2007; Boucher *et al*., 2016). In contrast, disordered sequences tend to become more structured even in the absence of selection. Notably, this directional change is not driven by biases in mutation probabilities arising from the genetic code or asymmetric nucleotide substitution rates, and is observed not only in random sequences but also in a large dataset of natural proteins, including conserved and *de novo* sequences. Together, these results suggest that the global sequence–structure landscape is diffuse, with structured and disordered proteins interspersed throughout sequence space (**??**). In contrast to fitness landscapes under selection, where valleys represent barriers to evolutionary trajectories, such barriers are largely absent under neutral evolution, making accessibility primarily a function of mutational connectivity. In such a landscape, mutations can move sequences in both more and less structured directions. However, because structured regions are distributed throughout sequence space, random mutational trajectories starting from disordered sequences are more likely, on average, to encounter increases in structure than decreases. This net directional effect would not arise if structured and disordered proteins were partitioned in distinct regions of the sequence space. While our model captures the general topology of the sequence–structure landscape, it does not preclude the existence of pronounced structural peaks or epistatic interactions. Instead, it suggests a landscape composed of dispersed hills of elevated structure embedded within a broadly disordered background. This interpretation is consistent with computational studies using simplified protein lattice models that support the existence of vast neutral networks, wherein similar structural phenotypes can be encoded by diverse genotypes (Bornberg-Bauer, 1997; Bastolla *et al*., 1999).

Our findings are unlikely to arise from artefacts of the disorder predictor used in the simulations (ADOPT). We obtained qualitatively similar results using IUPred3, which is based on a fundamentally different approach (Figure S2). While IUPred3 reproduced the relationship between the initial disorder score (*Z*) and its temporal change (quantified by *ρ*_*Zt*_), its lower resolution limited reliable fitting of the linear model.

Our results are also consistent with prior studies showing that structured conformations are broadly accessible in sequence space. For example, simulation-based work has shown that optimal structures for both proteins and RNAs can be reached from most starting configurations, owing to the rarity of fitness valleys — regions in which all neighboring configurations are less fit (Greenbury *et al*., 2022). Experimental studies provide similar evidence: highly active dihydrofolate reductase variants can be reached through only a few mutations despite a rugged fitness landscape (Papkou *et al*., 2023), and stable structures in the folding landscape of an anti-SARS-CoV-2 antibody remain accessible even in the presence of strong epistasis (Schulz *et al*., 2025). Notably, the latter study suggests that epistasis can facilitate, rather than hinder, accessibility. However, these studies primarily examine accessibility under selection. In contrast, our results show that structured states can become accessible even under neutrality, indicating that such accessibility is an intrinsic property of the sequence– structure landscape. This conclusion is further supported by prior work showing that secondary structure can emerge in long disordered regions during neutral evolution (Schaefer *et al*., 2010).

Our findings have several biological implications, particularly for the phenomenon of *de novo* gene emergence. Unlike genes derived from the divergence of existing protein-coding sequences, *de novo* genes are more likely to encode proteins that resemble random sequences (Middendorf *et al*., 2024). Empirical studies have shown that evolutionarily older *de novo* proteins tend to be less disordered than younger ones (Heames *et al*., 2020; Aubel *et al*., 2024). While this increase in structure could reflect selection for foldability and, by extension, physiological function, there is limited evidence directly linking structure to function in *de novo* proteins. Indeed, several *de novo* proteins with characterized functions are highly disordered (Aubel *et al*., 2023). Our model provides a potential explanation for these observations, showing that disordered sequences can acquire structure even in the absence of selection. This mechanism may therefore contribute to the structural evolution of *de novo* proteins.

In conclusion, our work elucidates a fundamental property of sequence–structure relationships and highlights how neutral evolution can be inherently constructive. It also opens avenues for targeted experimental validation using laboratory evolution approaches.

## Methods

### Generating random sequences

We generated random sequences using weighted sampling, parameterized by sequence length and amino acid frequencies (sampling weights). We described amino acid frequencies via GC-content, which influences codon probabilities and, in turn, amino acid composition.

We considered 30 parameter combinations defined by five sequence lengths (10, 30, 60, 90, and 120) and six GC-content values (30%, 35%, 40%, 45%, 50%, and 55%). We included five additional combinations using the same lengths but with uniform amino acid frequencies (1/20 for each amino acid).

For each of the 35 parameter combinations, we generated 10,000 random sequences. In all sequences, we fixed the first position to encode methionine.

### Protein evolutionary simulations

Our evolutionary simulations are based on the Wright–Fisher model with mutations. The process starts with a population of 100 identical sequences. At every generation, each sequence may undergo mutations, with the number of mutations per sequence drawn from a Poisson distribution with mean 1 (allowing zero mutations). Mutations occur uniformly along the sequence. The identity of each mutation is determined by an amino acid substitution matrix (*S*), where *S*_*ij*_ denotes the probability of substituting amino acid *i* with *j*.

We implemented the mutations via sequential random sampling. For example, if the Poisson draw equals 2, we sample two positions uniformly from the range [2, *L*], where *L* is the sequence length. This ensures that all positions except the first are allowed to mutate, while the first position remains fixed as methionine. If a selected position contains lysine, we use the corresponding row of the substitution matrix to obtain mutation probabilities, which are then used as weights to sample the substituted amino acid.

The next step simulates reproduction where the new generation completely replaces the current one (non-overlapping generations). We implement this by random sampling with replacement, assigning equal weights to all individuals to reflect neutral evolution.

We subjected each of the 35 × 10,000 random sequences to 30 generations of simulated evolution, and calculated structural disorder of the evolved sequences in each generation.

### Measuring structural disorder

To measure disorder of the protein sequences, we used ADOPT (Attention DisOrder Predic-Tor; Redl *et al*., 2023) that embeds protein sequences as numerical vectors using the ESM-1b protein language model and maps these embeddings to disorder scores via a regression model. The ADOPT output score (*Z*) is inversely related to disorder, with smaller *Z* values corresponding to higher disorder. We assigned one representative score per sequence by calculating the median of per-residue scores. We used the median because it reflects the typical level of disorder across the protein sequence and is less sensitive to extreme values than the mean. Specifically, it defines a threshold such that 50% of residues have higher scores. All mentions of the *Z* score in the manuscript refer to the median per-sequence score.

### Statistical analysis of disorder changes during evolution

From our preliminary correlation analysis, we found that the direction and magnitude of change in the *Z* score during evolution depend on the initial score (*Z*_0_). We used a linear model to test this observation more formally.

Specifically, we developed a linear model consistent with the Markov property of the evolutionary process (i.e., the population composition in the next generation depends only on that of the current generation). Formally, if *Z*_*t*_ denotes the mean *Z* score at generation *t* and Δ*Z*_*t*_ denotes the corresponding change (*Z*_*t*+1_ *− Z*_*t*_), the model is defined by Equation 1.

This formulation models the expected change in individual scores as a function of their current value, resulting in a corresponding population-level relationship in terms of mean *Z* scores.

Although this model does not explicitly include *Z*_0_, it captures its effect implicitly and offers several advantages over models that describe the full evolutionary trajectory:

- It aligns mechanistically with the simulation framework.
- It eliminates the need to track individual trajectories and their associated *Z*_0_ values.
- It increases statistical power by leveraging all time points within trajectories.

We fitted this model for each condition (length and composition) and tested the statistical significance of the fitted coefficients (*α* and *β*) using a t-test. Both coefficients were statistically significant in all cases (*P <* 10^*−*4^). We also calculated the pivot point (*Z*_*ϕ*_), defined as the value of *Z* for which the expected change (Δ*Z*) is zero, and estimated its 95% confidence interval using the delta method (Doob, 1935).

To assess the effect of sequence length, we fitted extended models that include length (*L*):

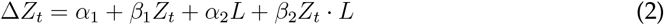

However, these models did not fit better than the individual models fitted separately for each condition, as the dependence of Δ*Z*_*t*_ on length was not monotonic.

### Randomizing mutation probabilities

We randomized amino acid substitution probabilities using two approaches. In the first approach, we randomized nucleotide substitution rates while keeping the genetic code structure intact. To this end, we randomly assigned probabilities to the six types of DNA substitutions (A:T→G:C, A:T→T:A, A:T→C:G, G:C→A:T, G:C→T:A and G:C→C:G) and computed the resulting amino acid substitution probabilities. In the second approach, we randomized the genetic code itself by randomly reshuffling the mapping between codons and amino acids while preserving the degeneracy structure of the standard genetic code. Thus, codons that encode the same amino acid in the standard genetic code (e.g. UUU and UUC) remain synonymous after reshuffling, although they may encode a different amino acid than in the canonical code (phenylalanine).

We applied these randomization procedures independently to each simulation (10,000 per condition), such that no two runs shared the same amino acid substitution probabilities.

## Notes

### Competing Interest Statement

The authors have declared no competing interest.

