## Supplementary material for "Structure can emerge from disorder under neutral evolution": Supplemetary Information

April 30, 2026

---

#### Contents

**1 Neutral mutations drive disordered proteins toward structure and structured proteins toward disorder** 2

**2 Directional disorder changes predicted using IUPred3 are qualitatively consistent with those obtained using ADOPT** 3

### 1 Neutral mutations drive disordered proteins toward structure and structured proteins toward disorder

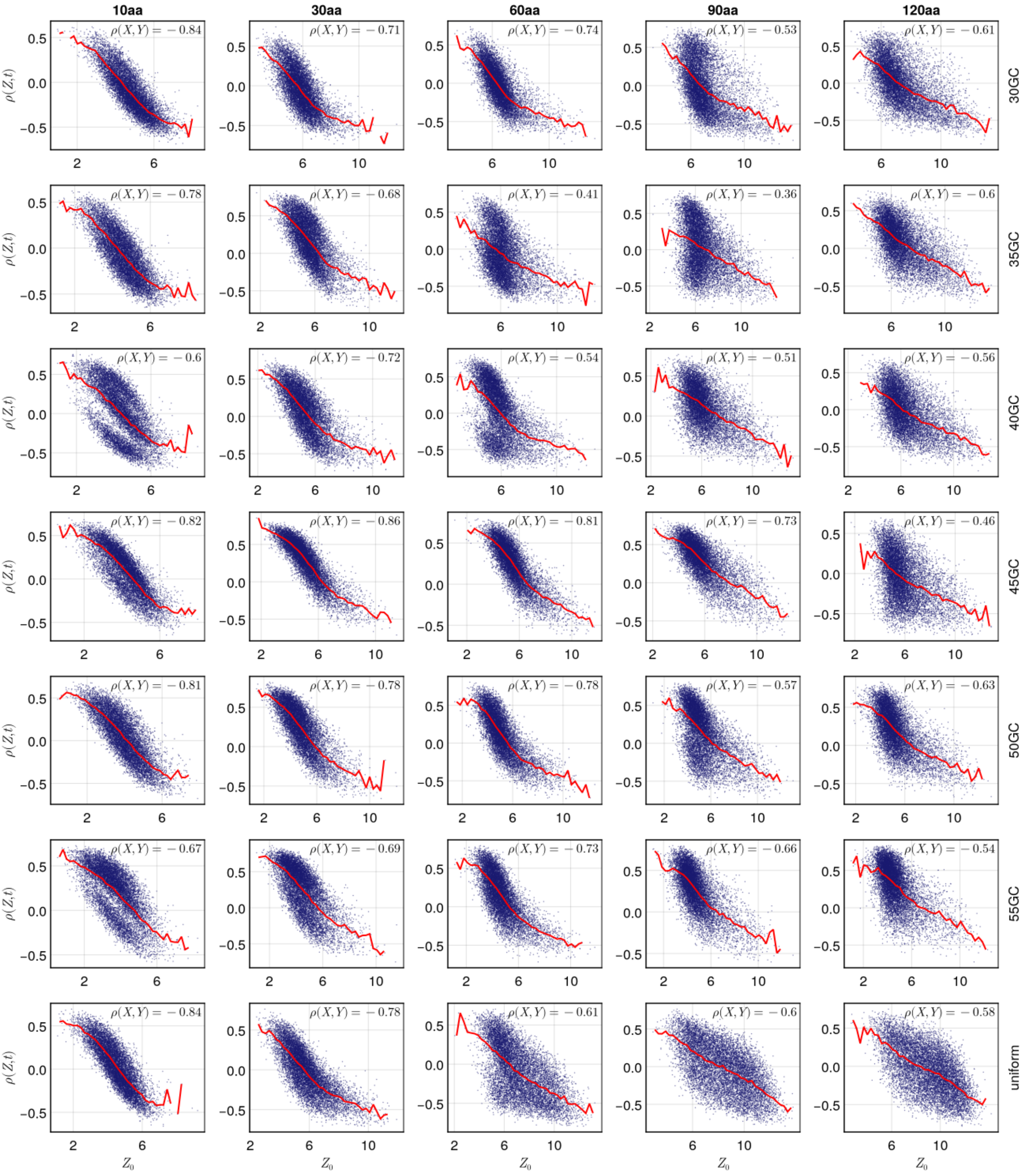


**Figure S1:** Scatter plots showing the relationship between temporal changes in $Z$ (quantified by $\rho_{Zt}$; vertical [X] axis) and the initial $Z$ score ($Z_0$; horizontal [Y] axis), for trajectories with sequence length 30 and uniform initial composition. The fitted trend line is shown in red. The inset reports the Spearman correlation between $X = \rho_{Zt}$ and $Y = Z_0$.

#### 2 Directional disorder changes predicted using IUPred3 are qualitatively consistent with those obtained using ADOPT

To ensure that our findings are not artefacts of the disorder predictor ADOPT, we repeated the analysis using IUPred3. Specifically, we randomly selected 50 trajectories per condition, excluding sequences of length 10, as this length smaller than the minimum length threshold of IUPred3. For each trajectory, we computed IUPred3 disorder scores for both the initial and evolved sequences.

IUPred3 disorder scores ($I$) were significantly correlated with ADOPT $Z$-scores (Spearman rank correlation: $-0.6 < \rho < -0.13$, $P < 10^{-99}$). The negative correlation is expected, as higher $I$-scores correspond to greater disorder, whereas higher $Z$-scores indicate lower disorder.

Next, we analysed how disorder change, quantified by the correlation between $I$-scores and time ($\rho_{It}$), depends on the initial disorder ($I_0$). Consistent with our original observations, $\rho_{It}$ decreases with increasing $I_0$ and transitions from positive to negative values at intermediate $I_0$. Although these relationships are noisier than those obtained using $Z$-scores, the qualitative trend is preserved. This increased variability is likely due to the smaller sample size (50 trajectories) compared to the original analysis (10,000 trajectories).

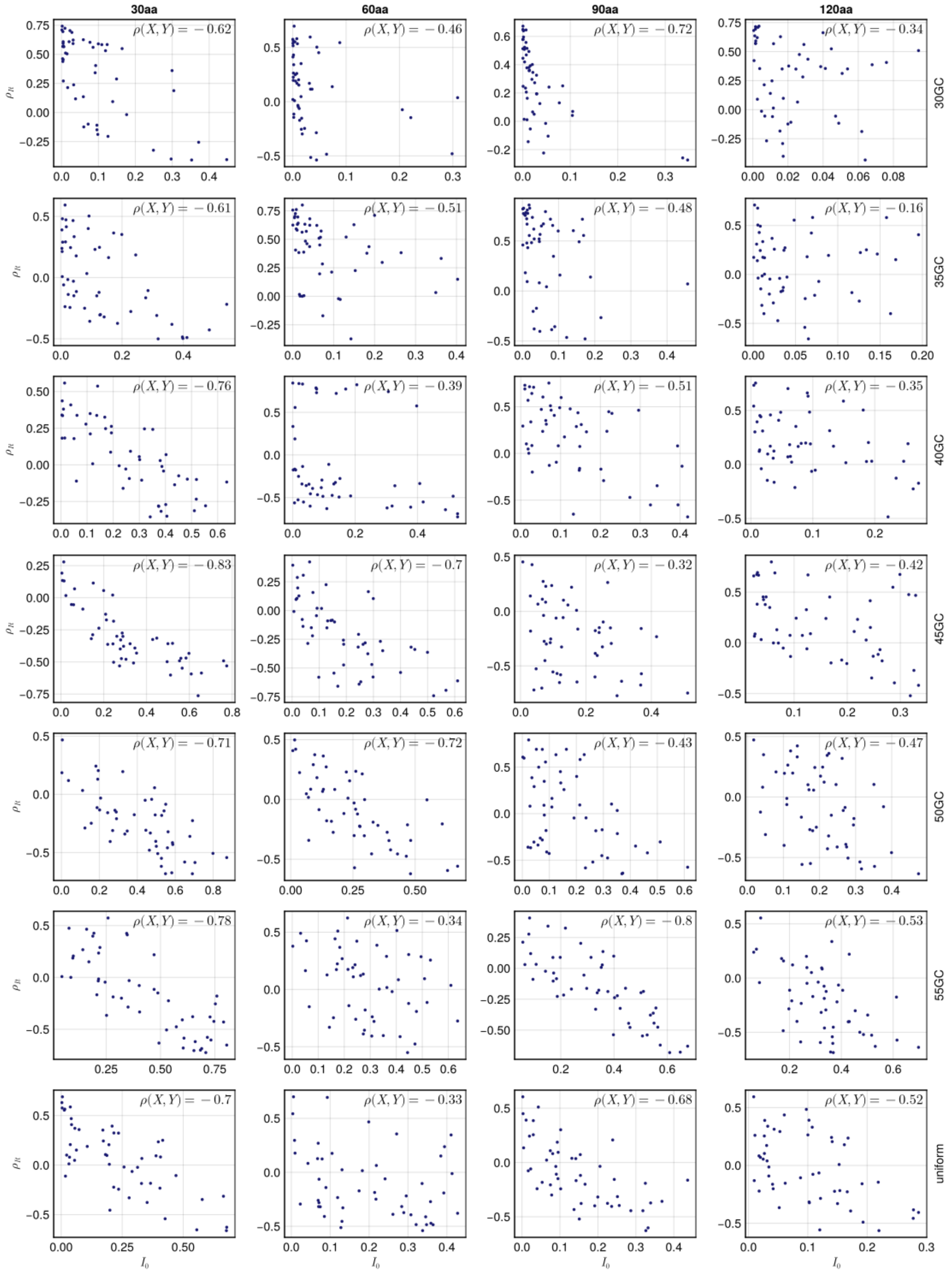


**Figure S2:** Scatter plots showing the relationship between temporal changes in $I$-scores (quantified by $\rho_{It}$; vertical [X] axis) and the initial $I$ score ($I_0$; horizontal [Y] axis), for trajectories with sequence length 30 and uniform initial composition. The inset reports the Spearman correlation between $X = \rho_{It}$ and $Y = I_0$.
